# A Comprehensive UHPLC-MS/MS and Extraction Method for Spinach (*Spinacia oleracea*) Flavonoids

**DOI:** 10.1101/2024.09.13.612955

**Authors:** Michael P. Dzakovich, Elaine A. Le, Alvin L. Tak, Shaji K. Chacko

## Abstract

Spinach produces an array of unique flavonoids not commonly found in other fruits and vegetables. These molecules likely serve as defense agents against biotic and abiotic stress and may have health beneficial properties for humans. Current methods to analyze spinach flavonoids are incomplete and only capture a portion of this uncharacterized pathway. A comprehensive analysis method is needed to determine how genetics, environmental conditions, and other factors influence spinach flavonoid biosynthesis. We developed and validated a high-throughput extraction and ultra high-performance liquid chromatography tandem mass spectrometry (UHPLC-MS/MS) method to separate and quantify 39 spinach flavonoid species in 11.5 minutes. Spinach flavonoids without authentic standards were putatively identified using MS/MS fragmentation experiments, precursor scans, and matches to high-resolution MS literature reports. Our extraction method enables up to 48 samples to be extracted in 60 minutes with recovery estimates between 100.5 – 107.8%. To assess the suitability of our method and generate benchmark estimates for 39 spinach flavonoids, we grew a panel of 30 genetically diverse spinach accessions and compared quantification data generated with a traditional or our high-throughput approach. Data generated by either approach were comparable, estimating total flavonoid averages of 75.1 – 170.1 or 93.1 – 187.26 mg/100 g fresh weight for the high-throughput and traditional method, respectively. Many estimates generated by our analysis method represent the first quantitative literature reports of these compounds. These experiments indicate that our extraction and analysis method is efficient, robust, and an important tool needed to study the biosynthesis and biological role of spinach flavonoids.

## INTRODUCTION

Flavonoids are secondary metabolites contributing to the pigmentation of flowers (aiding in pollination), quenching UV light in leaves, and protection against environmental biotic and abiotic stress (Takahashi and Ohnishi, 2004; Griesbach, 2005). Spinach (*Spinacia oleracea*) produces a remarkable concentration of flavonoids with varying reports between 82 – 224 mg/100 g fresh weight (Gil et al., 1999; Pandjaitan et al., 2005). Spinach-derived flavonoids, with reported antioxidant and anti-inflammatory properties, have been associated with a reduced risk of developing certain cancers, neurodegenerative, and cardiovascular diseases (Lomnitski et al., 2000; Edenharder et al., 2001; Nyska et al., 2001; Takahashi and Ohnishi, 2004; Grosso et al., 2017; Singh et al., 2018b; Hamsalakshmi et al., 2022). As research surrounding spinach flavonoids and their effects on human health continues to garner interest, lack of a comprehensive extraction and analysis methods represents a significant gap in knowledge. Moreover, the concentration range of spinach flavonoids remains unclear as many studies only focus on a few varieties of spinach and a few common flavonoids.

Spinach flavonoids are typically extracted by first homogenizing fresh or frozen spinach in a blender or mortar and pestle, followed by the addition of varying amounts of polar solvents including acetone, ethanol, water, and methanol, with the latter two being used most frequently (Edenharder et al., 2001; Koh et al., 2012; Singh et al., 2017). Samples are then centrifuged and flavonoids can be quantified from filtered supernatants (Cho et al., 2008; Koh et al., 2012; Singh et al., 2017, 2018a, 2018b). Current extraction methods tend to be time-consuming, and more efficient methods are needed to profile larger populations. Quantification of spinach flavonoids is generally done by reverse phase liquid chromatography tandem mass spectrometry (LC-MS/MS), although other less sensitive approaches have also been implemented such as photodiode array detector (Howard et al., 2002; Pandjaitan et al., 2005; Singh et al., 2018b). Through ^1^H and ^13^C NMR, as well as mass spectrometry, over 40 flavonoid species have been identified in spinach (Ferreres et al., 1997; Singh et al., 2018b). However, most published methods only consider 5-10 of these flavonoids with the most comprehensive method quantifying up to 17 (Koh et al., 2012). Despite a lack of authentic standards for most of these compounds, common product ions have been determined (e.g. 330 *m*/*z*) for many of these flavonoids using accurate mass collision-induced dissociation (CID) experiments (Singh et al., 2019). These data make it possible to create a comprehensive analysis method that simultaneously quantifies most of the known flavonoids in spinach. Utilizing an efficient extraction technique with a comprehensive quantification method would enable scientists to determine the diversity of flavonoids in spinach frequently utilized by consumers. These data could better inform investigators seeking to determine the role of spinach flavonoids in human health.

To gain additional insight into the diversity of spinach flavonoids, we developed a high-throughput extraction (48 samples/hour) and comprehensive quantification method (39 flavonoids/11.5 minutes) using high-performance liquid chromatography tandem mass spectrometry (UHPLC-MS/MS). In addition to traditional validation experiments, we profiled a panel of 30 genetically diverse spinach accessions using both a validated and high-throughput extraction method to determine how flavonoid estimates and population structures differed due to methodology. Here, we summarize this series of experiments that validated our methods and provide benchmark concentrations for spinach flavonoids in a genetically diverse population.

## MATERIALS AND METHODS

### Chemical and Reagents

Methanol (MeOH; LC-MS grade), water (LC-MS grade), formic acid (FA; LC-MS grade), and taxifolin (99.99%) were purchased from Thermo Fisher Scientific (Pittsburgh, PA, United States). Naringin (≥95% purity), naringenin (≥95%), and quercetin-3-glucoside (≥95%) were purchased from Sigma-Aldrich (St. Louis, MO, United States). Spinacetin (>95%) and jaceidin (>95%) were purchased from Key Organics (Camelford, United Kingdom). Patuletin (≥98%) was purchased from Extrasynthese (Genay, France). Acetonitrile (LC-MS grade) was purchased from VWR (Radnor, PA, United States).

### Standard Curve and Internal Standard Preparation

Unless stated otherwise, all solutions were prepared in 1:1 MeOH:H_2_O + 0.1% FA to maximize analyte solubility and chromatographic resolution. Stock solutions of each authentic standard were made by dissolving weighed powders in MeOH + 0.1% FA and diluting 1:1 with H_2_O + 0.1% FA to final concentrations of 0.1 mg/mL. Thirty μL of each stock solution was combined into a 4 mL glass vial and 36 μL of 20 μM taxifolin was added (4% of final volume). This concentration was selected as it generates a signal intensity that is intermediate to the range of intensities seen for all analytes quantified by our method. An additional 714 μL of 1:1 MeOH:H_2_O + 0.1% FA was added to bring the final volume of the stock solution to 900 μL. Eight additional 4 mL vials with 600 μL of 1:1 MeOH:H_2_O + 0.1% FA were made and a 3x dilution series was created spanning to 6561x dilution. Depending on the analyte, concentrations ranged from 9.03 nmol/vial to 0.63 pmol/vial. Due to the unavailability of glycosylated spinach flavonoid standards, these analytes were quantified relative to quercetin-3-glucoside (Singh et al., 2017; Grace et al., 2022)

Solutions of 200 nmol naringin and naringenin were also prepared for spike recovery experiments and used to routinely monitor extraction efficiency for each sample. These molecules are not detectable in spinach but have similar chromatographic characteristics as glycosylated and aglycone spinach flavonoids, respectively.

### Sample Material

To maximize genetic variation, 30 spinach accessions previously phenotyped and genotyped were selected and grown (Qin et al., 2017; Hayes et al., 2020). Selections were made based on reported differences in carotenoid and mineral concentrations as well as genetic features that segregated accessions into unique subgroups. Metadata about each accession can be found in Supplemental Table 1. Before sowing, seeds were osmoprimed in tap water and stored in a 4 refrigerator for five days to enhance germination rates. Osmoprimed seeds were sown into one-gallon pots filled with HFC/20 Growing mix (Jolly Gardener Products; Oviedo, FL) and thinned to six plants per pot. Plants were grown in a PGW36 walk-in growth chamber (Conviron; Winnipeg, Canada) maintained at 20 ± 0.5 °C / 15 ± 0.5 °C day and night temperatures, respectively, with a 12 hour photoperiod (300 μmol/m^2^/s of photosynthetically active radiation) and pot positions were randomized on a daily basis. Relative humidity was maintained at 50 ± 10%. Plants were irrigated with deionized water until germination and then sub-irrigated with a nutrient solution containing: 1.2 mM KNO_3_, 0.8 mM Ca(NO_3_)_2_, 0.8 mM NH_4_NO_3_, 0.2 mM MgSO_4_, 0.3 mM KH_2_PO_4_, 25 μM CaCl_2_, 25 μM H_3_BO_3_, 2 μm MnSO_4_, 2 μM ZnSO_4_, 0.5 μM CuSO_4_, 0.5 μM H_2_MoO_4_, 0.1 μM NiSO_4_, and 10 μM Fe (III)-N, N′-ethylenebis[2-(2-hydroxyphenyl)-glycine] (Sprint 138; Becker-Underwood Inc.; Ames, IA) according to Qin and colleagues (Qin et al., 2017). Plants were harvested when each plant within a pot had six to eight fully developed leaves (approximately 35 days after sowing). Tissue was frozen at −80 °C. Frozen samples were homogenized in a ratio of 1:1 spinach:MilliQ H_2_O with a VWR Model 250 polytron at 20,000 RPM for ∼one minute. Homogenate was aliquoted and immediately stored at −80 °C until extraction and analysis.

### Quality Control (QC) Material and Pseudo-Standard Preparation

A quality control (QC) composite sample was created by blending equal parts of the 30 accessions referenced above into a single homogenate. This homogenate was aliquoted for spike recovery experiments, intra/interday variation experiments, and used for MS/MS validation studies described below.

A methanolic extract of the QC material was diluted 1:1 with H_2_O + 0.1% FA and semi-purified using a solid phase extraction (SPE; StrataX 33 μM, 30 mg, 3 mL tube; Phenomenex, Torrance, CA, United States) method adapted from Redan and others (Redan et al., 2017). Spinach flavonoids were eluted from cartridges using 500 μL of MeOH + 0.1% FA and the eluent was diluted 1:1 with H_2_O + 0.1% FA. These samples were used for multiple reaction monitoring (MRM) optimization experiments.

### Flavonoid Extraction

#### Traditional method

Spinach samples (500 mg ± 10 mg) were extracted using a validated extraction protocol (Mohamedshah et al., 2022). Samples were spiked (200 μL) with a 1:1 solution of 200 nmol naringin and 200 nmol naringenin in 1:1 MeOH:H_2_O + 0.1% FA. Samples were extracted with 5 mL of 98:2 MeOH:FA, vortexed for 1 minute, sonicated for 20 minutes, and centrifuged for 5 minutes at 3700 RPM at 4°C in a VWR Mega Star 4.0R. The supernatant was decanted into 15 mL borosilicate glass tubes. This extraction process was repeated twice more but sonicated for only 10 minutes.

#### High throughput method

A high-throughput extraction using a Tecan Freedom EVO 150 (Tecan; Mannedorf, Switzerland) was adapted from previous methodology validated for carotenoids and modified for water soluble flavonoids (Dzakovich et al., 2023). Samples (100 ± 5 mg) were spiked with 100 μL of 1:1 200 nmol naringin to 200 nmol naringenin internal standard solution. Two stainless steel balls (Grainger #4RJH5, l2 inch diameter) were then inserted into each sample tube. After several minutes to allow internal standards to equilibrate with sample matrix, 1 mL of 98:2 MeOH:FA was added to each sample. Samples were then extracted in a Spex SamplePrep 2010 Geno/Grinder for 45 seconds at 1400 RPM and centrifuged for 3 minutes at 13,000 RPM. The supernatant was collected into 15 mL borosilicate glass tubes and the extraction process was repeated twice more.

#### Drying, resolubilizing, and filtering

For both extraction methods, samples were loaded into a Labconco RapidVap (Kansas City, MO, United States) with lid heater for drying. While maintaining a temperature of 35 and 35% speed, the vacuum was gradually decreased in 20 mbar increments from a starting pressure of 280 mbar to 10 mbar. After drying, samples were capped and stored at −80 °C until quantification. For samples extracted using the traditional method, 5 mL of 1:1 MeOH:H_2_O + 0.1% FA was added to each tube and sonicated (∼15 seconds) with agitation to resuspend dried residue on tube walls. Samples were vortexed and a 1 mL aliquot was removed and diluted in 4 mL of 1:1 MeOH:H_2_O + 0.1% FA with internal standard (3800 μL of 1:1 MeOH:H_2_O + 0.1% FA + 200 μL of 20 μM taxifolin in the same carrier solvent) in a separate tube. This dilution step ensured analyte concentrations were within linear detection limits of the instrument. For the high-throughput extraction, 200 μL of 20 μM taxifolin solution was spiked into each tube prior to adding 4800 μL of 1:1 MeOH:H_2_O + 0.1% FA. Samples were then sonicated as described above. For both extraction methods, redissolved samples were vortexed and filtered through a 0.2 μm PTFE syringe filter into HPLC vials and run immediately.

### Flavonoid Quantification

Spinach flavonoids were separated on a Waters C_18_ Acquity bridged ethylene hybrid (BEH) 2.1x100 mm, 1.7 mm particle size column installed in a ThermoFisher Vanquish Horizon UHPLC with a column chamber maintained at 40 interfaced with a ThermoFisher TSQ Altis triple quadrupole mass spectrometer. The autosampler was maintained at 12 to minimize sample oxidation. A 0.5 mL/min gradient was developed consisting of mobile phase A (LC-MS grade water + 0.1% FA) and mobile phase B (LC-MS grade acetonitrile + 0.1% FA) were adjusted as follows: 0% B to 6.0% B over 0.5 minutes, 6.0% B to 9.0% B for 1.5 minutes, 9% B to 13% B over 1.0 minute, 13% B to 35% B over 1.5 minutes, 35% B to 75% B over 2.91 minutes, 75% B to 90% B over 0.09 minutes, 90% B held for 2.0 minutes, 90% B to 0% B over 0.5 minutes, hold at 0% B for an additional 1.5 minutes to equilibrate the column. Each run was a total of 11.5 minutes and the needle was washed twice for 5 seconds using MeOH + 0.1% FA followed by water. To minimize matrix buildup on the mass spectrometer sampling cone and optics, column eluent was routed to waste for the first 1.9 minutes and the last 4.0 minutes of each run utilizing a post-column diverter valve. Mass spectrometer source and MRM parameters can be found in Table 1. Dwell times were calculated for each analyte to ensure 12-15 points per peak. Since glycosylated spinach flavonoids are not commercially available, quantification was relative to quercetin-3-glucoside using the linear portion of a 9-point standard curve. The aglycones jaceidin, patuletin, and spinacetin were quantified against commercially available authentic standards. Peak areas were normalized to taxifolin spiked into samples and standards to correct for within- and between-day instrument variability.

**Table 1.**
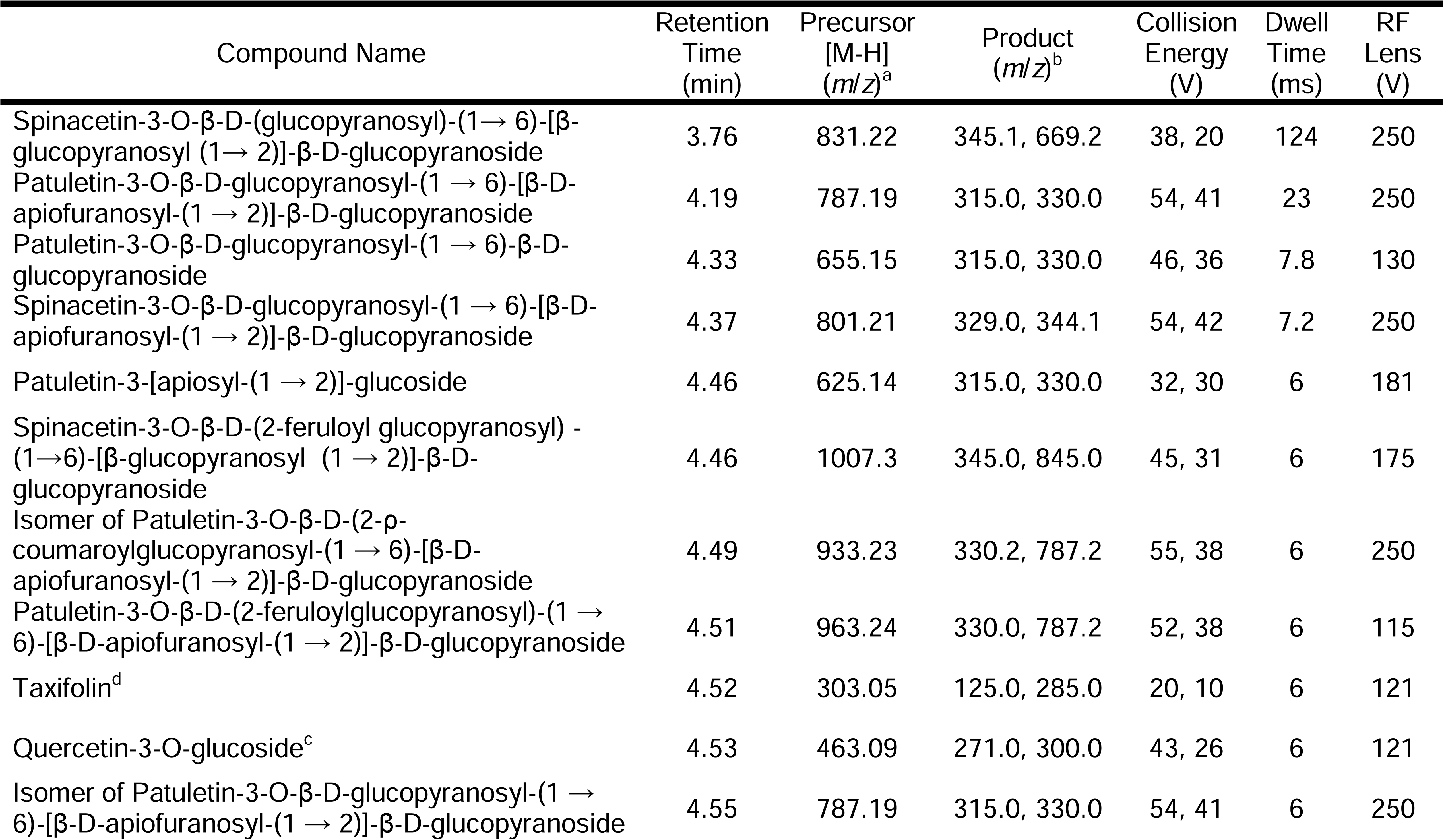
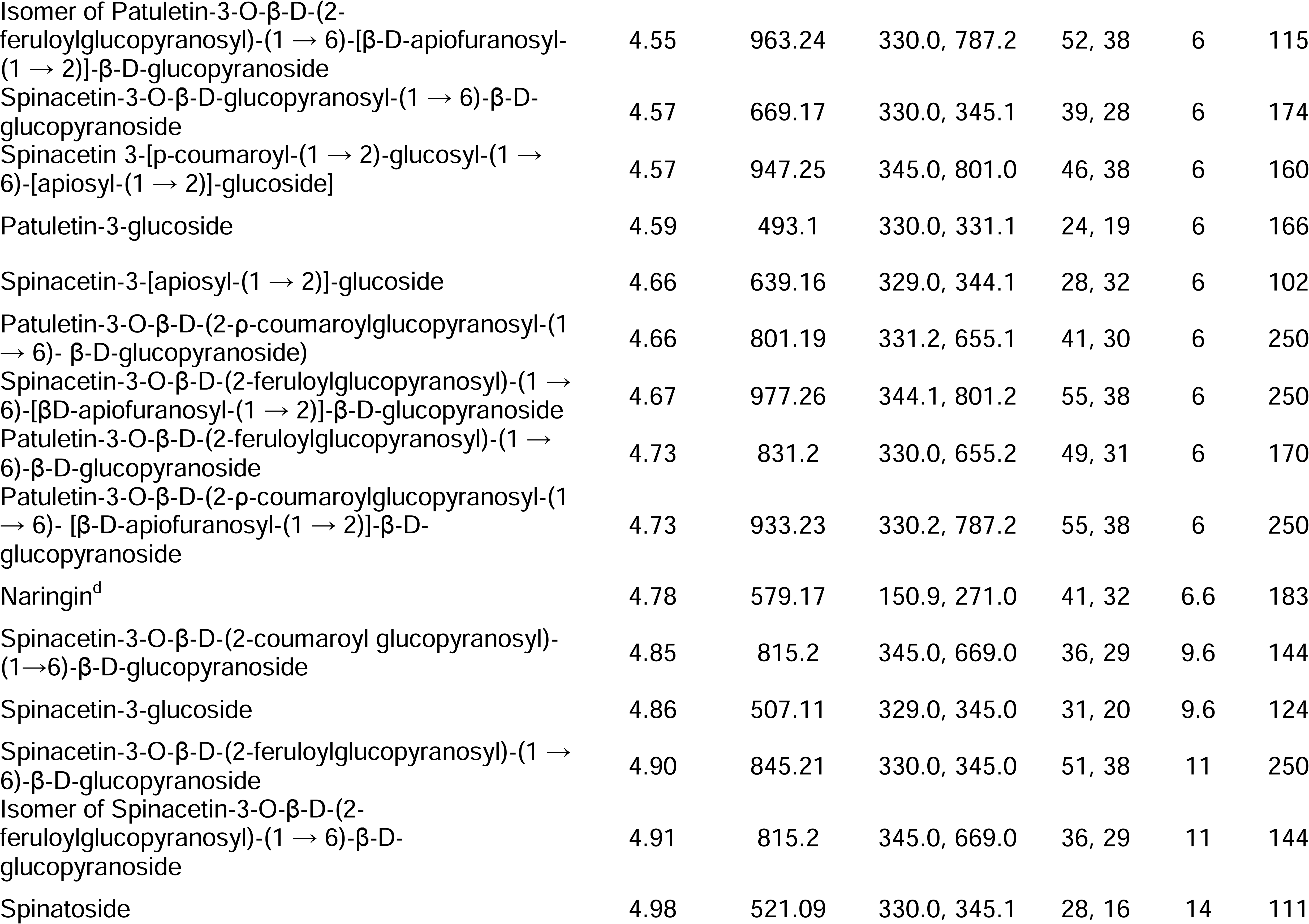
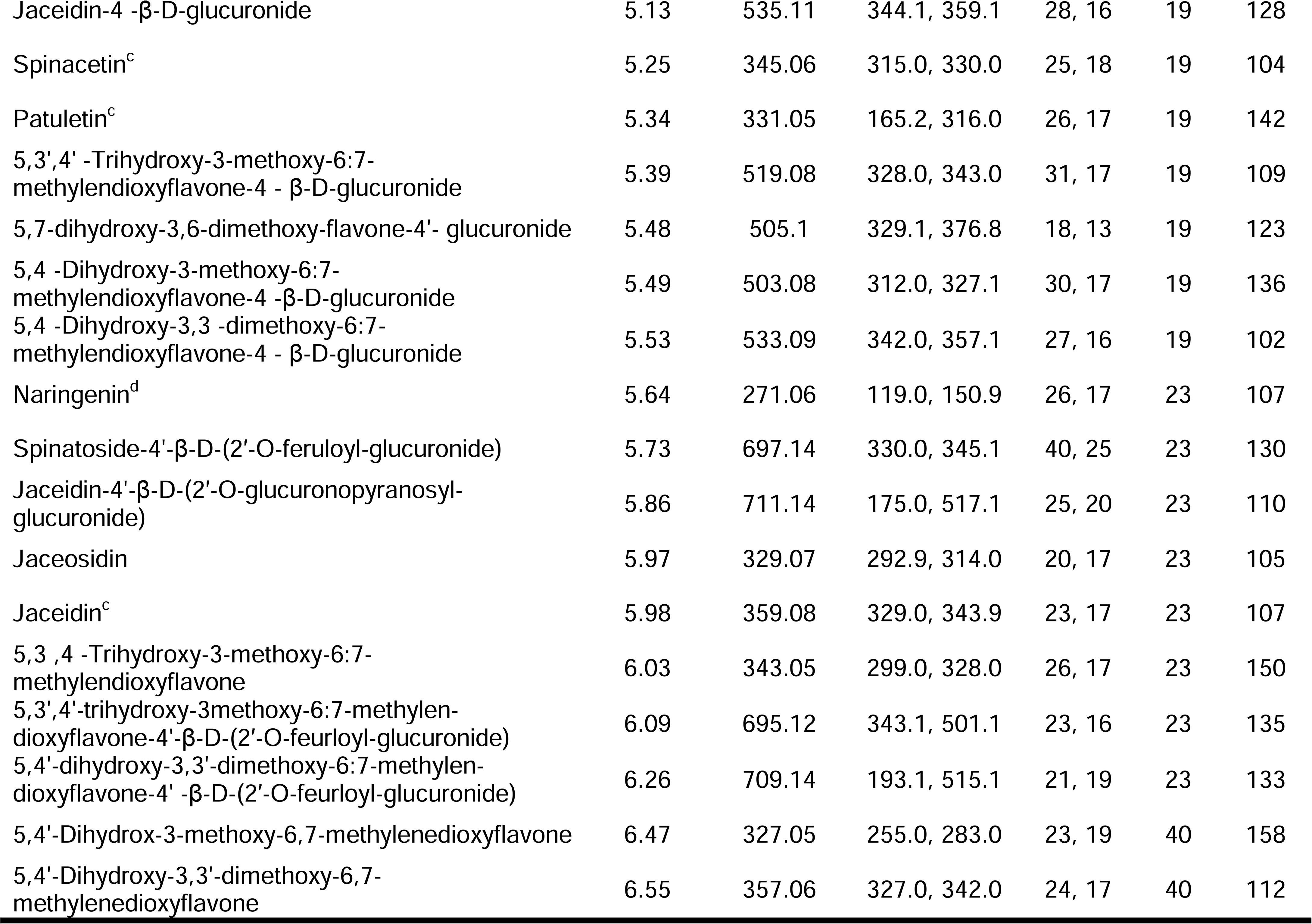
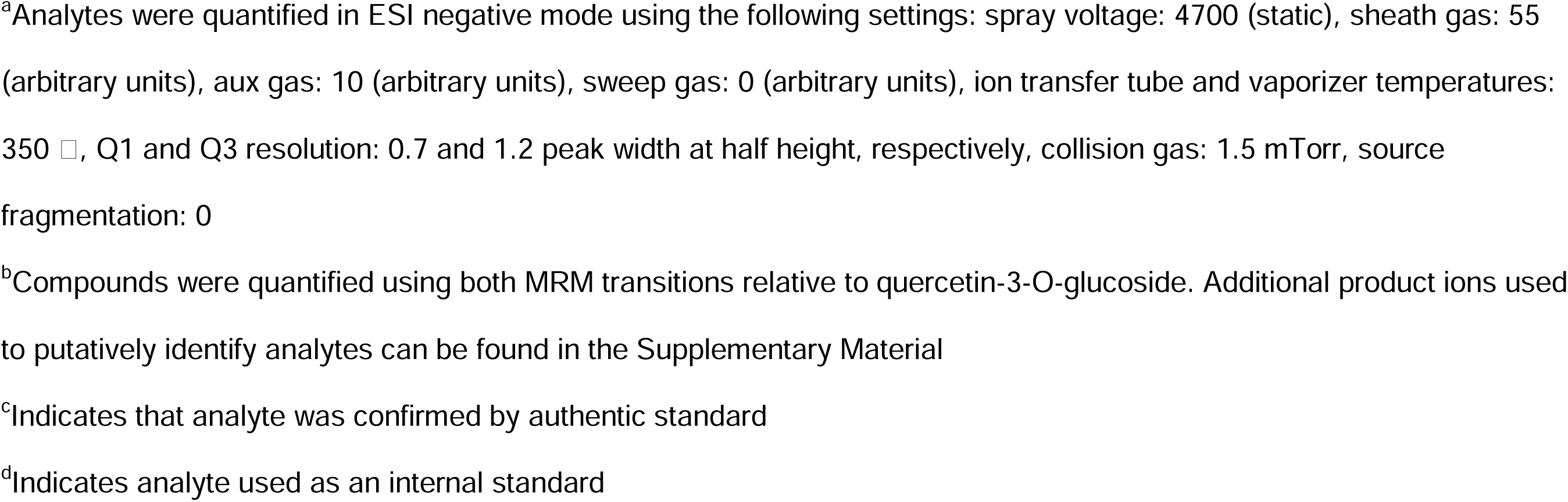
LC-MS/MS MRM parameters of quantified spinach flavonoids and internal/external standards.

| Compound Name | Retention Time (min) | Precursor [M-H] (*m/z*)<sup>a</sup> | Product (*m/z*)<sup>b</sup> | Collision Energy (V) | Dwell Time (ms) | RF Lens (V) |
| --- | --- | --- | --- | --- | --- | --- |
| Spinacetin-3-O-β-D-(glucopyranosyl)-(1 → 6)-[β-glucopyranosyl (1 → 2)]-β-D-glucopyranoside | 3.76 | 831.22 | 345.1, 669.2 | 38, 20 | 124 | 250 |
| Patuletin-3-O-β-D-glucopyranosyl-(1 → 6)-[β-D-apiofuranosyl-(1 → 2)]-β-D-glucopyranoside | 4.19 | 787.19 | 315.0, 330.0 | 54, 41 | 23 | 250 |
| Patuletin-3-O-β-D-glucopyranosyl-(1 → 6)-β-D-glucopyranoside | 4.33 | 655.15 | 315.0, 330.0 | 46, 36 | 7.8 | 130 |
| Spinacetin-3-O-β-D-glucopyranosyl-(1 → 6)-[β-D-apiofuranosyl-(1 → 2)]-β-D-glucopyranoside | 4.37 | 801.21 | 329.0, 344.1 | 54, 42 | 7.2 | 250 |
| Patuletin-3-[apiosyl-(1 → 2)]-glucoside | 4.46 | 625.14 | 315.0, 330.0 | 32, 30 | 6 | 181 |
| Spinacetin-3-O-β-D-(2-feruloyl glucopyranosyl) - (1 → 6)-[β-glucopyranosyl (1 → 2)]-β-D-glucopyranoside | 4.46 | 1007.3 | 345.0, 845.0 | 45, 31 | 6 | 175 |
| Isomer of Patuletin-3-O-β-D-(2-p-coumaroylglucopyranosyl-(1 → 6)-[β-D-apiofuranosyl-(1 → 2)]-β-D-glucopyranoside | 4.49 | 933.23 | 330.2, 787.2 | 55, 38 | 6 | 250 |
| Patuletin-3-O-β-D-(2-feruloylglucopyranosyl)-(1 → 6)-[β-D-apiofuranosyl-(1 → 2)]-β-D-glucopyranoside | 4.51 | 963.24 | 330.0, 787.2 | 52, 38 | 6 | 115 |
| Taxifolin<sup>d</sup> | 4.52 | 303.05 | 125.0, 285.0 | 20, 10 | 6 | 121 |
| Quercetin-3-O-glucoside<sup>c</sup> | 4.53 | 463.09 | 271.0, 300.0 | 43, 26 | 6 | 121 |
| Isomer of Patuletin-3-O-β-D-glucopyranosyl-(1 → 6)-[β-D-apiofuranosyl-(1 → 2)]-β-D-glucopyranoside | 4.55 | 787.19 | 315.0, 330.0 | 54, 41 | 6 | 250 |
| Isomer of Patuletin-3-O-β-D-(2-feruloylglucopyranosyl)-(1 → 6)-[β-D-apiofuranosyl-(1 → 2)]-β-D-glucopyranoside | 4.55 | 963.24 | 330.0, 787.2 | 52, 38 | 6 | 115 |
| Spinacetin-3-O-β-D-glucopyranosyl-(1 → 6)-β-D-glucopyranoside | 4.57 | 669.17 | 330.0, 345.1 | 39, 28 | 6 | 174 |
| Spinacetin 3-[p-coumaroyl-(1 → 2)-glucosyl-(1 → 6)-[apiosyl-(1 → 2)]-glucoside] | 4.57 | 947.25 | 345.0, 801.0 | 46, 38 | 6 | 160 |
| Patuletin-3-glucoside | 4.59 | 493.1 | 330.0, 331.1 | 24, 19 | 6 | 166 |
| Spinacetin-3-[apiosyl-(1 → 2)]-glucoside | 4.66 | 639.16 | 329.0, 344.1 | 28, 32 | 6 | 102 |
| Patuletin-3-O-β-D-(2-p-coumaroylglucopyranosyl)-(1 → 6)- β-D-glucopyranoside) | 4.66 | 801.19 | 331.2, 655.1 | 41, 30 | 6 | 250 |
| Spinacetin-3-O-β-D-(2-feruloylglucopyranosyl)-(1 → 6)-[β-D-apiofuranosyl-(1 → 2)]-β-D-glucopyranoside | 4.67 | 977.26 | 344.1, 801.2 | 55, 38 | 6 | 250 |
| Patuletin-3-O-β-D-(2-feruloylglucopyranosyl)-(1 → 6)-β-D-glucopyranoside | 4.73 | 831.2 | 330.0, 655.2 | 49, 31 | 6 | 170 |
| Patuletin-3-O-β-D-(2-p-coumaroylglucopyranosyl)-(1 → 6)- [β-D-apiofuranosyl-(1 → 2)]-β-D-glucopyranoside | 4.73 | 933.23 | 330.2, 787.2 | 55, 38 | 6 | 250 |
| Naringin<sup>d</sup> | 4.78 | 579.17 | 150.9, 271.0 | 41, 32 | 6.6 | 183 |
| Spinacetin-3-O-β-D-(2-coumaroyl glucopyranosyl)-(1→6)-β-D-glucopyranoside | 4.85 | 815.2 | 345.0, 669.0 | 36, 29 | 9.6 | 144 |
| Spinacetin-3-glucoside | 4.86 | 507.11 | 329.0, 345.0 | 31, 20 | 9.6 | 124 |
| Spinacetin-3-O-β-D-(2-feruloylglucopyranosyl)-(1 → 6)-β-D-glucopyranoside | 4.90 | 845.21 | 330.0, 345.0 | 51, 38 | 11 | 250 |
| Isomer of Spinacetin-3-O-β-D-(2-feruloylglucopyranosyl)-(1 → 6)-β-D-glucopyranoside | 4.91 | 815.2 | 345.0, 669.0 | 36, 29 | 11 | 144 |
| Spinatoside | 4.98 | 521.09 | 330.0, 345.1 | 28, 16 | 14 | 111 |
| Jaceidin-4 -β-D-glucuronide | 5.13 | 535.11 | 344.1, 359.1 | 28, 16 | 19 | 128 |
| Spinacetin<sup>c</sup> | 5.25 | 345.06 | 315.0, 330.0 | 25, 18 | 19 | 104 |
| Patuletin<sup>c</sup> | 5.34 | 331.05 | 165.2, 316.0 | 26, 17 | 19 | 142 |
| 5,3',4' -Trihydroxy-3-methoxy-6:7-methylendioxyflavone-4 - β-D-glucuronide | 5.39 | 519.08 | 328.0, 343.0 | 31, 17 | 19 | 109 |
| 5,7-dihydroxy-3,6-dimethoxy-flavone-4' - glucuronide | 5.48 | 505.1 | 329.1, 376.8 | 18, 13 | 19 | 123 |
| 5,4 -Dihydroxy-3-methoxy-6:7-methylendioxyflavone-4 -β-D-glucuronide | 5.49 | 503.08 | 312.0, 327.1 | 30, 17 | 19 | 136 |
| 5,4 -Dihydroxy-3,3 -dimethoxy-6:7-methylendioxyflavone-4 - β-D-glucuronide | 5.53 | 533.09 | 342.0, 357.1 | 27, 16 | 19 | 102 |
| Naringenin<sup>d</sup> | 5.64 | 271.06 | 119.0, 150.9 | 26, 17 | 23 | 107 |
| Spinatoside-4'-β-D-(2'-O-feruloyl-glucuronide) | 5.73 | 697.14 | 330.0, 345.1 | 40, 25 | 23 | 130 |
| Jaceidin-4'-β-D-(2'-O-glucuronopyranosyl-glucuronide) | 5.86 | 711.14 | 175.0, 517.1 | 25, 20 | 23 | 110 |
| Jaceosidin | 5.97 | 329.07 | 292.9, 314.0 | 20, 17 | 23 | 105 |
| Jaceidin<sup>c</sup> | 5.98 | 359.08 | 329.0, 343.9 | 23, 17 | 23 | 107 |
| 5,3 ,4 -Trihydroxy-3-methoxy-6:7-methylendioxyflavone | 6.03 | 343.05 | 299.0, 328.0 | 26, 17 | 23 | 150 |
| 5,3',4'-trihydroxy-3methoxy-6:7-methylen-dioxyflavone-4'-β-D-(2'-O-feurloyl-glucuronide) | 6.09 | 695.12 | 343.1, 501.1 | 23, 16 | 23 | 135 |
| 5,4'-dihydroxy-3,3'-dimethoxy-6:7-methylen-dioxyflavone-4' -β-D-(2'-O-feurloyl-glucuronide) | 6.26 | 709.14 | 193.1, 515.1 | 21, 19 | 23 | 133 |
| 5,4'-Dihydrox-3-methoxy-6,7-methylenedioxyflavone | 6.47 | 327.05 | 255.0, 283.0 | 23, 19 | 40 | 158 |
| 5,4'-Dihydroxy-3,3'-dimethoxy-6,7-methylenedioxyflavone | 6.55 | 357.06 | 327.0, 342.0 | 24, 17 | 40 | 112 |

<sup>a</sup>Analytes were quantified in ESI negative mode using the following settings: spray voltage: 4700 (static), sheath gas: 55 (arbitrary units), aux gas: 10 (arbitrary units), sweep gas: 0 (arbitrary units), ion transfer tube and vaporizer temperatures: 350 °C, Q1 and Q3 resolution: 0.7 and 1.2 peak width at half height, respectively, collision gas: 1.5 mTorr, source fragmentation: 0
<sup>b</sup>Compounds were quantified using both MRM transitions relative to quercetin-3-O-glucoside. Additional product ions used to putatively identify analytes can be found in the Supplementary Material
<sup>c</sup>Indicates that analyte was confirmed by authentic standard
<sup>d</sup>Indicates analyte used as an internal standard

### Limit of Detection and Quantification

Six replicates of the lowest concentration calibrant from standard curves for quercetin-3-glucoside, patuletin, spinacetin, and jaceidin were used to calculate limit of detection (LOD; 3:1 signal to noise) and limit of quantification (LOQ; 10:1 signal to noise) (Vial and Jardy, 1999). A weighting factor for the signal to noise for each analyte’s lowest standard curve calibrant was calculated to estimate moles on column at 10:1 or 3:1 signal to noise.

### Spike addition experiments

Quality control samples (100 mg ± 5 mg) were randomly divided into two different groups of six tubes, 1) tubes only containing spinach homogenate; 2) tubes containing spinach homogenate spiked with standards. Additionally, another 6 tubes only spiked with standards were also utilized. 20 μL spikes consisted of: 200 nmol naringin, 200 nmol naringenin, and 50 nmol jaceidin. Samples underwent a traditional extraction and were analyzed using the LC-MS/MS method described above. Spike recovery was calculated using the following formula:

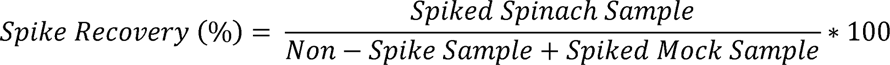

### Intra/Interday Variability Experiments

The intra/interday variability experiments compared the quantification values from six replicates from the same quality control material over three days. Each day, a set of 6 randomly selected samples (100 mg ± 5 mg) in 2 mL tubes was spiked with 100 μL of 1:1 200 nM naringin: 200 nM naringenin, accompanied by three internal standard-only tubes. The samples then underwent a high-throughput extraction. Intra and interday variability was calculated by determining the coefficient of variation for each analyte within a day or between days, respectively.

### Random Effects Modeling and Data Visualization

Random effects modeling and data visualization were conducted using R v4.4.1 (R Development Core Team, 2018). Random effects models were run to partition variance associated with genetic background, extraction type, and replicate using the package lme4 (Bates et al., 2015). The following model was used:

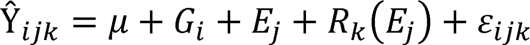

In the above model, Ŷ*_ijk_* represents an analyte estimate within the *i*th genotype, *j*th extraction method, and *k*th replicate; μ represents the population mean of an analyte; *G_i_* represents the contribution from genetic factors for the *i*th genotype of the population; *E_j_* represents the contribution from extraction for the *j*th extraction method; *R_k_*(*E_J_*) represents the contribution of the *k*th replicate within extraction method and can be interpreted as within-method variation; and ε*_ijk_* represents the residual error.

Coordinates for 2-D scores plots were generated using the packages FactoMineR and Factoextra, scaling variable means to zero and standard deviation one, as input data for principal components analysis (PCA) (Lê et al., 2008). ggplot2 and MoMA color palette were used for visualization (Wickham, 2016; Paull et al., 2024).

## RESULTS AND DISCUSSION

### Development and Validation of UHPLC-MS/MS Method

#### Precursor Ion Discovery

To maximize the detection, separation, and quantification of flavonoids that are unique to spinach, a combined literature search and precursor ion scan approach was utilized. Previous mass spectrometry based approaches revealed over 20 species of flavonoids present in spinach and provide valuable fragmentation information that can be used to putatively identify molecules in the absence of authentic standards (Cho et al., 2008). For example, most of the flavonoids identified by Cho and colleagues generated either a 333 or 347 *m*/*z* ion in positive mode. High-resolution mass spectrometry enabled the discovery of glucuronidated flavonoids in spinach as well as several other glycosylated molecules (Singh et al., 2017, 2018a, 2019). Merging the findings of the last 20 years of method development yields a list of 39 flavonoid molecules discovered in spinach with varying degrees of information available about them (Supplemental Table 2). We utilized this information to determine if each of these compounds could be detected in methanolic spinach extracts.

We created a methanolic extract of the quality control material described above and injected it into a ThermoFisher Vanquish UHPLC system in tandem with a TSQ Altis triple quadrupole mass spectrometer operated in negative mode. Using a gradient developed for anthocyanins (Mengist et al., 2020), we separated our extract on a Waters Acquity BEH column (2.1 x 100 mm; 1.7 μm particle size). We developed selected ion recordings (SIRs) for each candidate flavonoid. Additionally, we ran precursor ion scans by leveraging fragmentation information available about common core structures found in most spinach flavonoids (Fig. 1) to eliminate isobaric peaks that are not flavonoids. We explored a variety of collision energies to find precursor ions that produced product ions consistent with patuletin (e.g. 316 *m*/*z*), spinacetin (e.g. 330 *m*/*z*), and jaceidin (e.g. 344 *m*/*z*) as well as other product ions produced by carbon monoxide/dioxide loses, M-122 losses, demethylation, and retro-Diels-Alder reactions (Fabre et al., 2001; McNab et al., 2009; Dimkić et al., 2020). Unoptimized MRM scans helped further reduce extraneous peaks using previously validated product ions (Singh et al., 2018a, 2019; Grace et al., 2022). Product ion scans of the remaining chemical features, their relative retention order, and matches to database/literature reference helped putatively identify the remaining chemical features where commercial standards are not available. A summary of this information can be found in Supplemental Table 2.

**Fig. 1.**
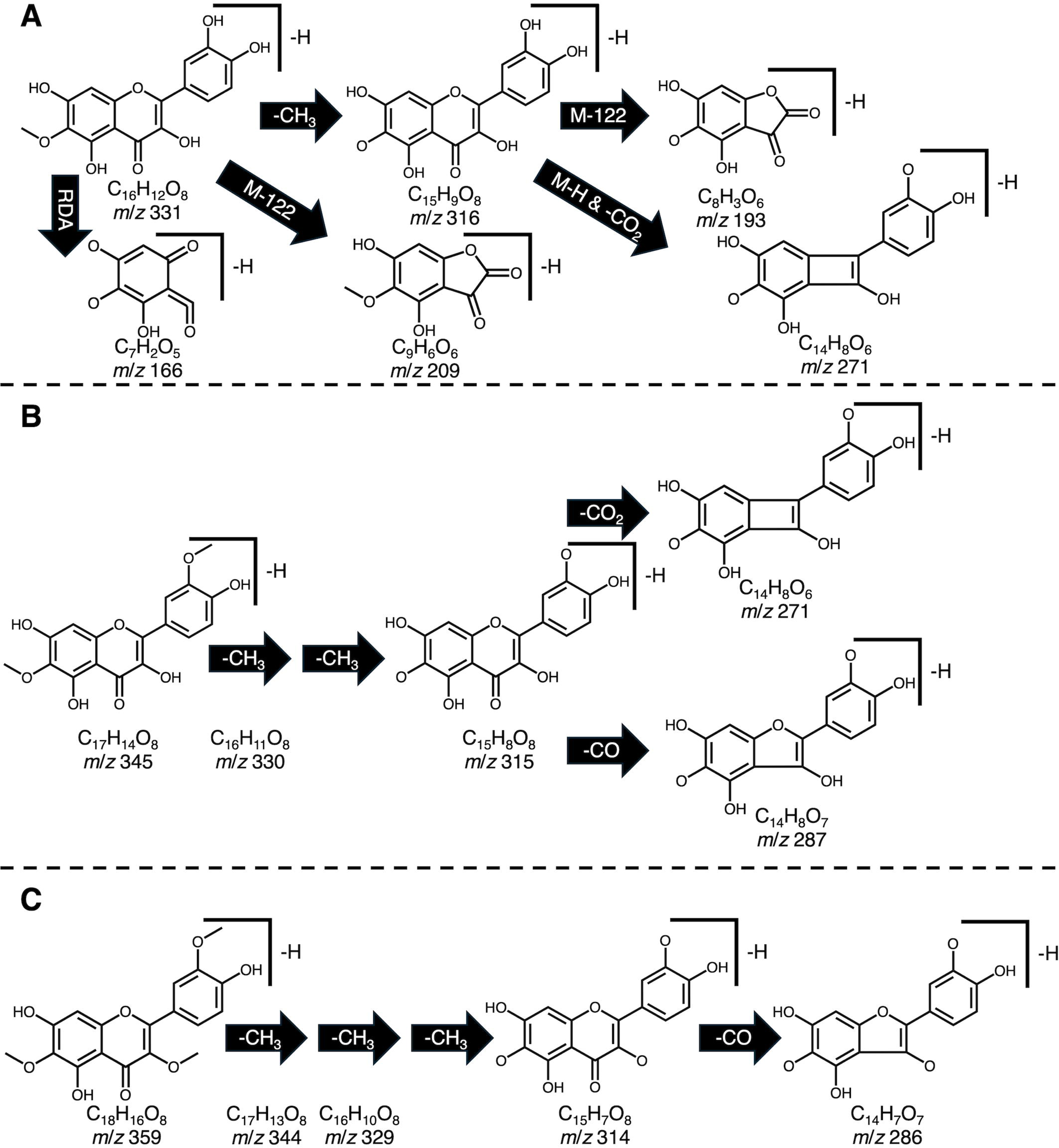
Example fragmentation patterns reported for patuletin (A), spinacetin (B), and jaceidin (C) including proton loss (M-H), retro Diels-Alder (RDA), demethylation (-CH_3_), carbon monoxide loss (-CO), carbon dioxide loss (-CO_2_). Fragmentation patterns were used to guide identification efforts to screen for potential precursor molecules containing these motifs.

#### Internal Standard Selection

We explored five different internal standard candidates commonly used in flavonoid analysis including naringenin, naringin, ethyl gallate, phlorizin, and taxifolin (Mengist et al., 2020; Grace et al., 2022; Mohamedshah et al., 2022). Ethyl gallate and phlorizin were eliminated due to their structural dissimilarity to spinach flavonoids, stability, and ionization efficiency. Spikes of 200 μM naringin/naringenin (200 or 100 μL depending on extraction method) generated peak areas consistent with those seen in spinach flavonoids quantified by our method, eluted near analytes, and are not naturally present in spinach in detectable amounts.

#### Optimization of MS parameters

Source temperature parameters (ion transfer tube and vaporizer temperatures) were optimized by successively injecting a mixture of quercetin-3-glucoside, naringin, naringenin, spinacetin, jaceidin, and patuletin into the TSQ Altis at temperatures ranging from 275 to 375 at 25 intervals. At a flow rate of 0.5 mL/min, 350 for both source parameters resulted in the highest average signal intensity for these analytes.

Positive and negative ionization modes were explored since existing mass spectrometry approaches for spinach flavonoids have reported both (Cho et al., 2008; Singh et al., 2017, 2018a). We consistently found that while our analytes ionized in both modes, negative mode tended to result in higher signal to noise ratios compared to positive mode. Moreover, there is a greater abundance of fragmentation information for spinach flavonoids in negative mode allowing for more confident identification (Singh et al., 2017, 2018a, 2019).

To optimize MRMs, semi-purified flavonoid extracts described earlier were infused into the TSQ Altis source. The flow rate of the UHPLC was maintained at 0.5 mL/min and eluent composition was adjusted to match the ratio of mobile phases where each analyte was determined to elute. The TSQ Altis systematically tested spray voltages (1500 – 5000 V), sheath gas (0 – 60 arbitrary units), auxiliary gas (0 – 25 arbitrary units), sweep gas (0 – 20 arbitrary units), RF lens voltage (30 – 250 V), and collision energy (5 – 55 V) seeking five product ions per analyte. For all parameters except RF lens voltage and collision energy, average values were determined to provide the most parsimonious setpoint for all analytes. The top two MRMs with the highest signal to noise ratio were retained for quantification and other pertinent MS information can be found in Table 1. Additional qualifying product ions for each analyte as well as structures can be found in the Supplemental Information (Supplemental Table 2). Windows of 15 seconds on each side of a retention time were added to maximize the instrument’s duty cycle and allow for 12-15 points across a peak. Analytes available as authentic standards were optimized as outlined above but without SPE.

#### Chromatographic gradient

We initially used a chromatographic gradient developed for separating anthocyanins using both Waters BEH and HSS T3 2.1 x 100 mm (1.7μm particle size) C_18_ columns (Mengist et al., 2020). Generally, the BEH column exhibited more favorable separation of relatively polar spinach flavonoids compared to the HSS T3. The original gradient used 1% FA as a modifier for their mobile phase A to aid in pH maintenance, chromatography, and ionization nuances important for anthocyanins. We found that switching to 0.1% FA for both mobile phases slightly improved signal of most analytes, ostensibly from a reduction in formate adducts, and lowers the overall cost of mobile phase preparation. To shorten the overall runtime of our method to 11.5 minutes, we initiated a jump to 90% B (acetonitrile + 0.1% FA) shortly after our final analyte eluted followed by an equilibration period allowing for the column to be stabilized by the following injection cycle. A two-phase wash cycle of MeOH + 0.1% FA followed by H_2_O + 0.1% FA ensured minimal carryover. Multiple analytes in our method exhibited isomers that have been previously reported in the literature (Cho et al., 2008; Singh et al., 2017). Each of these isomers were chromatographically resolved from their respective counterparts allowing for separate quantification. All 39 spinach flavonoids analyzed by our method are separated in under 3 minutes with the remaining time devoted to column cleaning and equilibration.

#### LOD/LOQ

UV-Vis spectral information for many spinach flavonoids have been reported (Cho et al., 2008; Singh et al., 2018a; Grace et al., 2022), but overwhelmingly similar spectral characteristics make it impossible to distinguish many spinach flavonoids from one another. Mass spectrometry allows for the development of highly selective and sensitive methods while overcoming chromatography issues such as co-elution. Limits of detection and quantification do not appear to be published for previous methods developed for these compounds, so we sought to calculate these values here. Our method was able to detect jaceidin, patuletin, spinacetin, and quercetin-3-glucoside at sub-femtomole-on-column quantities (Table 2). Assuming the same extraction, redissolve, and injection conditions, we estimate that flavonoids at 23.6 pmol/gram spinach tissue or higher could be quantified. Sensitivity at this level enables high-throughput, micro-extraction methods where little sample (< 100 mg) is required.

**Table 2.** Extraction efficiency of commercially available spinach flavonoids and structurally similar internal standards.

| Analyte | Extraction Efficiency (%) | LOQ (femtomoles injected) | LOD (femtomoles injected) | LOQ (picomoles per sample)<sup>c</sup> | LOD (picomoles per sample) |
| --- | --- | --- | --- | --- | --- |
| Jaceidin | 107.8±6.3 | 0.193 | 0.058 | 0.966 | 0.289 |
| Patuletin<sup>a</sup> | NA | 1.078 | 0.323 | 5.389 | 1.617 |
| Spinacetin<sup>a</sup> | NA | 0.369 | 0.111 | 1.849 | 0.555 |
| Quercetin-3-glucoside<sup>a</sup> | NA | 1.193 | 0.358 | 5.964 | 1.789 |
| Naringin<sup>b</sup> | 102.1±2.1 | NA | NA | NA | NA |
| Naringenin<sup>b</sup> | 100.5±4.7 | NA | NA | NA | NA |

<sup>a</sup>Analyte used solely for quantification <sup>b</sup>Analyte used as an internal standard with no calibration curve <sup>c</sup>Estimate assumes a 5 µL injection, 25 mL redissolve volume, and 0.5 mg sample of 1:1 spinach:water homogenate

### Validation of Extraction

Given the speed of our chromatographic gradient, the possibility to merge our analysis method with a high-throughput extraction method poses an exciting opportunity to quantitatively profile large populations of spinach in a relatively short period of time.

Miniaturized extractions are quickly gaining popularity as they reduce solvent usage, less labor-intensive, and are often amenable to automation compared to traditional methods. We developed a high-throughput protocol based on previously validated approaches for vegetables and cereals (Dzakovich et al., 2020, 2023) making modifications for spinach. We conducted spike recovery and intra/interday variability experiments to determine the suitability of our high-throughput approach. Additionally, we compared both the traditional extraction method with our high-throughput approach by profiling a genetically diverse population of spinach. Published spinach flavonoid values are sparse and infrequently use multiple genotypes (Howard et al., 2002; Pandjaitan et al., 2005; Cho et al., 2008).

#### Spike Recovery

A spike addition experiment allowed us to generate quantitative recovery estimates from our high-throughput method. We excluded spinacetin and patuletin from this experiment because these aglycones are rarely present in detectable levels in spinach. Naringin (102.1 ± 2.1%) and naringenin (100.5 ± 4.7%) had recovery estimates similar to jaceidin (107.8 ± 6.3%), which is naturally present in our sample matrix (Table 2). These findings support that our high-throughput extraction works efficiently with analytes varying in polarity. Our recovery estimates mirror or exceed estimates for similar molecules (Furrer et al., 2017; Mengist et al., 2020; Mohamedshah et al., 2022).

#### Intra/Interday variability

Inter and intraday variability experiments allowed us to quantify the combined variation of our high-throughput extraction method as well as our analysis. A single individual extracted six randomly selected samples and analyzed them using our UHPLC-MS/MS method. This experiment was repeated twice more by the same individual to determine interday variation. Our findings indicate that our high-throughput extraction can reliably estimate analyte concentrations both within and between days (Table 3). The use of taxifolin as an internal standard helped compensate for day-day variation that is common in mass spectrometers.

**Table 3.**
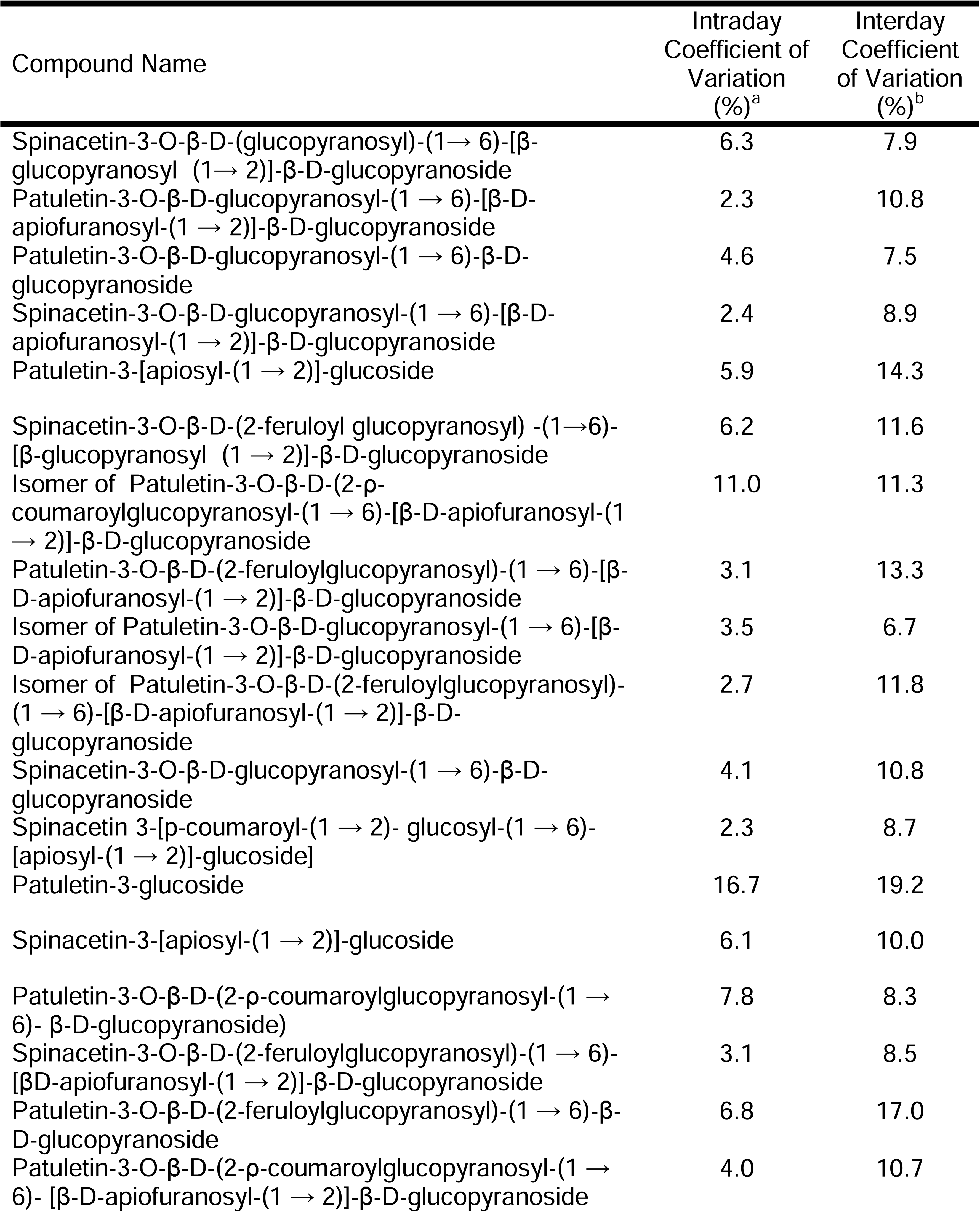
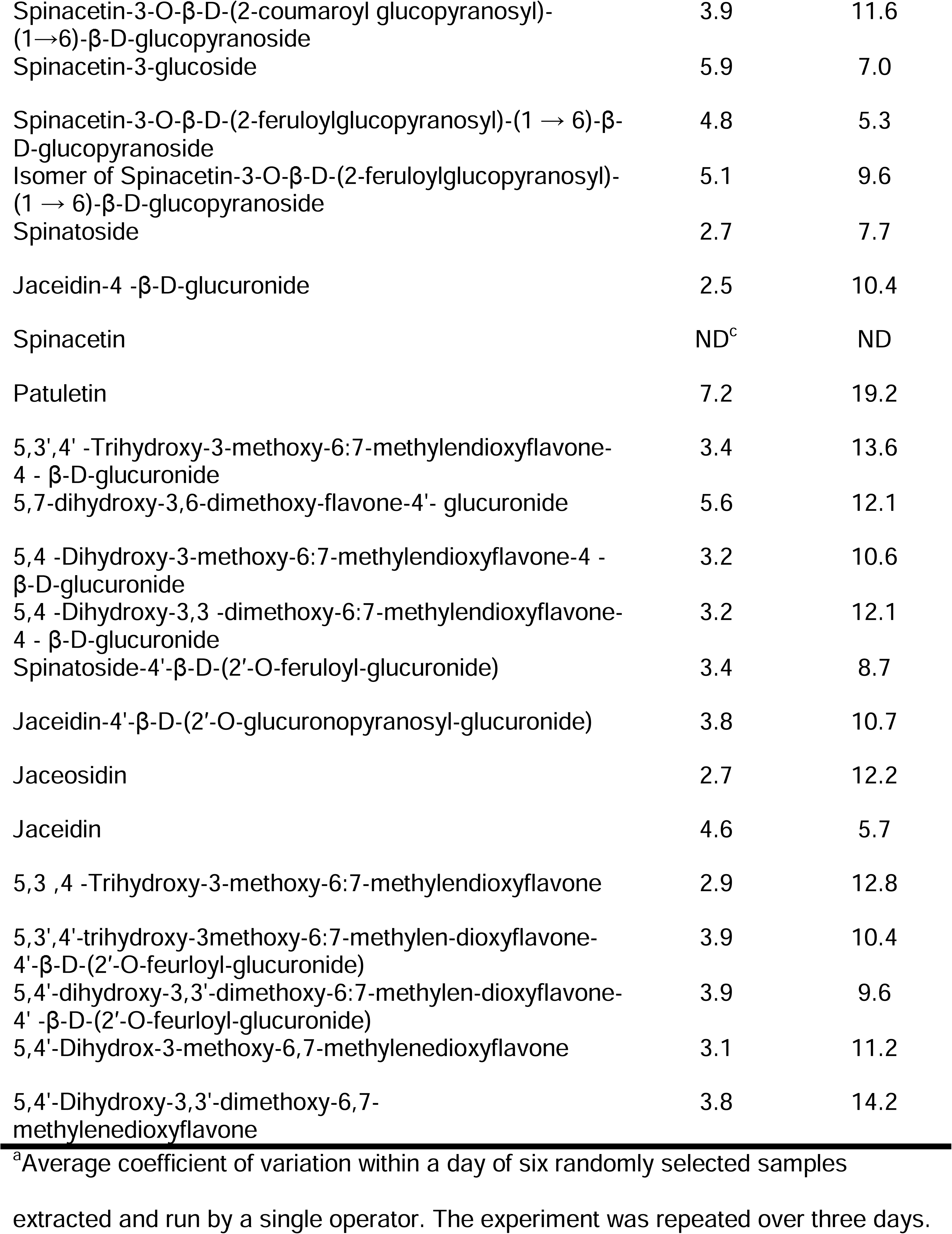
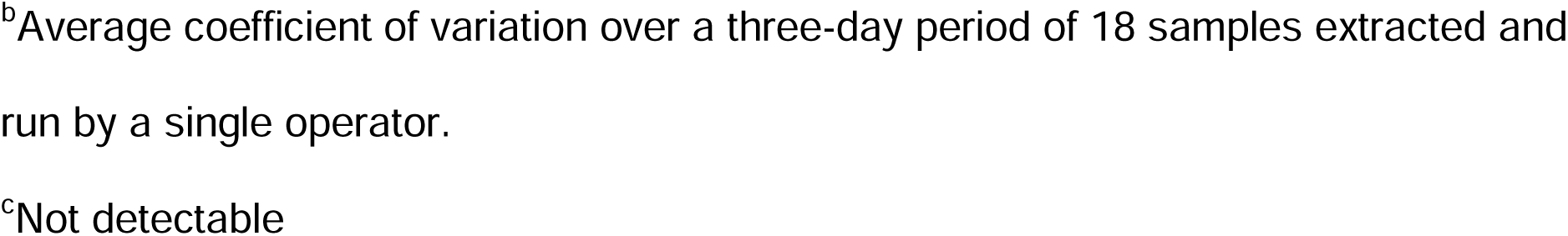
Intraday and interday coefficient of variation values for analytes separated and quantified by our UHPLC-MS/MS method extracted using our high-throughput method.

| Compound Name | Intraday<br>Coefficient of<br>Variation<br>(%)<sup>a</sup> | Interday<br>Coefficient<br>of Variation<br>(%)<sup>b</sup> |
| --- | --- | --- |
| Spinacetin-3-O-β-D-(glucopyranosyl)-(1→6)-[β-glucopyranosyl (1→2)]-β-D-glucopyranoside | 6.3 | 7.9 |
| Patuletin-3-O-β-D-glucopyranosyl-(1→6)-[β-D-apiofuranosyl-(1→2)]-β-D-glucopyranoside | 2.3 | 10.8 |
| Patuletin-3-O-β-D-glucopyranosyl-(1→6)-β-D-glucopyranoside | 4.6 | 7.5 |
| Spinacetin-3-O-β-D-glucopyranosyl-(1→6)-[β-D-apiofuranosyl-(1→2)]-β-D-glucopyranoside | 2.4 | 8.9 |
| Patuletin-3-[apiosyl-(1→2)]-glucoside | 5.9 | 14.3 |
| Spinacetin-3-O-β-D-(2-feruloyl glucopyranosyl) -(1→6)-[β-glucopyranosyl (1→2)]-β-D-glucopyranoside | 6.2 | 11.6 |
| Isomer of Patuletin-3-O-β-D-(2-p-coumaroylglucopyranosyl-(1→6)-[β-D-apiofuranosyl-(1→2)]-β-D-glucopyranoside | 11.0 | 11.3 |
| Patuletin-3-O-β-D-(2-feruloylglucopyranosyl)-(1→6)-[β-D-apiofuranosyl-(1→2)]-β-D-glucopyranoside | 3.1 | 13.3 |
| Isomer of Patuletin-3-O-β-D-glucopyranosyl-(1→6)-[β-D-apiofuranosyl-(1→2)]-β-D-glucopyranoside | 3.5 | 6.7 |
| Isomer of Patuletin-3-O-β-D-(2-feruloylglucopyranosyl)-(1→6)-[β-D-apiofuranosyl-(1→2)]-β-D-glucopyranoside | 2.7 | 11.8 |
| Spinacetin-3-O-β-D-glucopyranosyl-(1→6)-β-D-glucopyranoside | 4.1 | 10.8 |
| Spinacetin 3-[p-coumaroyl-(1→2)- glucosyl-(1→6)-[apiosyl-(1→2)]-glucoside] | 2.3 | 8.7 |
| Patuletin-3-glucoside | 16.7 | 19.2 |
| Spinacetin-3-[apiosyl-(1→2)]-glucoside | 6.1 | 10.0 |
| Patuletin-3-O-β-D-(2-p-coumaroylglucopyranosyl-(1→6)- β-D-glucopyranoside) | 7.8 | 8.3 |
| Spinacetin-3-O-β-D-(2-feruloylglucopyranosyl)-(1→6)-[βD-apiofuranosyl-(1→2)]-β-D-glucopyranoside | 3.1 | 8.5 |
| Patuletin-3-O-β-D-(2-feruloylglucopyranosyl)-(1→6)-β-D-glucopyranoside | 6.8 | 17.0 |
| Patuletin-3-O-β-D-(2-p-coumaroylglucopyranosyl-(1→6)- [β-D-apiofuranosyl-(1→2)]-β-D-glucopyranoside | 4.0 | 10.7 |
| Spinacetin-3-O-β-D-(2-coumaroyl glucopyranosyl)-<br>(1→6)-β-D-glucopyranoside | 3.9 | 11.6 |
| Spinacetin-3-glucoside | 5.9 | 7.0 |
| Spinacetin-3-O-β-D-(2-feruloylglucopyranosyl)-(1 → 6)-β-<br>D-glucopyranoside | 4.8 | 5.3 |
| Isomer of Spinacetin-3-O-β-D-(2-feruloylglucopyranosyl)-<br>(1 → 6)-β-D-glucopyranoside | 5.1 | 9.6 |
| Spinatoside | 2.7 | 7.7 |
| Jaceidin-4 -β-D-glucuronide | 2.5 | 10.4 |
| Spinacetin | ND<sup>c</sup> | ND |
| Patuletin | 7.2 | 19.2 |
| 5,3',4' -Trihydroxy-3-methoxy-6:7-methylendioxyflavone-4 - β-D-glucuronide | 3.4 | 13.6 |
| 5,7-dihydroxy-3,6-dimethoxy-flavone-4'- glucuronide | 5.6 | 12.1 |
| 5,4 -Dihydroxy-3-methoxy-6:7-methylendioxyflavone-4 -<br>β-D-glucuronide | 3.2 | 10.6 |
| 5,4 -Dihydroxy-3,3 -dimethoxy-6:7-methylendioxyflavone-4 - β-D-glucuronide | 3.2 | 12.1 |
| Spinatoside-4'-β-D-(2'-O-feruloyl-glucuronide) | 3.4 | 8.7 |
| Jaceidin-4'-β-D-(2'-O-glucuronopyranosyl-glucuronide) | 3.8 | 10.7 |
| Jaceosidin | 2.7 | 12.2 |
| Jaceidin | 4.6 | 5.7 |
| 5,3 ,4 -Trihydroxy-3-methoxy-6:7-methylendioxyflavone | 2.9 | 12.8 |
| 5,3',4'-trihydroxy-3methoxy-6:7-methylen-dioxyflavone-4'-β-D-(2'-O-feurloyl-glucuronide) | 3.9 | 10.4 |
| 5,4'-dihydroxy-3,3'-dimethoxy-6:7-methylen-dioxyflavone-4' -β-D-(2'-O-feurloyl-glucuronide) | 3.9 | 9.6 |
| 5,4'-Dihydrox-3-methoxy-6,7-methylenedioxyflavone | 3.1 | 11.2 |
| 5,4'-Dihydroxy-3,3'-dimethoxy-6,7-methylenedioxyflavone | 3.8 | 14.2 |

<sup>a</sup>Average coefficient of variation within a day of six randomly selected samples
extracted and run by a single operator. The experiment was repeated over three days.
<sup>b</sup>Average coefficient of variation over a three-day period of 18 samples extracted and run by a single operator.
<sup>c</sup>Not detectable

#### Spinach germplasm survey

To further test our high-throughput extraction and simultaneously address the lack of comprehensive, quantitative information on spinach flavonoids, we profiled a population of 30 spinach accessions. Accessions include both commonly grown/consumed spinach cultivars that are commercially available as well as wild-collected accessions curated by the USDA Germplasm Resources Information Network (USDA-GRIN). Details about the individual accessions as well as analyte concentrations generated by both traditional and high-throughput extractions can be found in the Supplemental Information (Supplemental Tables 1 and 3). Total flavonoid concentrations ranged from 75.1 – 170.1 mg/100 g fresh weight and approximately 70% of total flavonoids was represented by 519.08, 801.21, 521.09, 533.09, 787.19, and 977.26 *m*/*z* (Supplemental Tables 3 and 4). Using the traditional method, data ranged from 93.1 – 187.26 mg/100 g fresh weight. Both datasets align well with previous reports of flavonoid concentrations in spinach (Cho et al., 2008; Grace et al., 2022). Critically, the proportion of each flavonoid compared to total flavonoids in each method were nearly identical (Supplemental Table 4). This finding indicates that differences between extraction methods are largely due to differences in extraction efficiency. Our data additionally indicate that spinach produces substantially greater amounts of flavonoids compared to other fruits and vegetables (Harnly et al., 2006; Slimestad et al., 2008). Given the limited environmental variability in our study, the range of concentrations seen in our data primarily reflect genetic differences. Environmental aspects such as light and temperature could drastically alter the concentrations of spinach flavonoids (Howard et al., 2002; Cho et al., 2008). While overall trends in data were similar between extraction methods (Supplemental Tables 3 and 4), some differences exist.

Principal components analysis visualized the population structure as a function of extraction and illustrates that differences within and among accessions was slightly less pronounced in the high-throughput data (Figure 2B). Notably, replicates of the same accession were further apart, indicating less similarity. A larger portion of variance explained by principal component 1 allowed for a clearer delineation among different accessions in the traditionally extracted samples. In practice, population structures were nearly identical and an end-user such as a plant breeder would be able to generalize trends using either method.

**Fig. 2.**
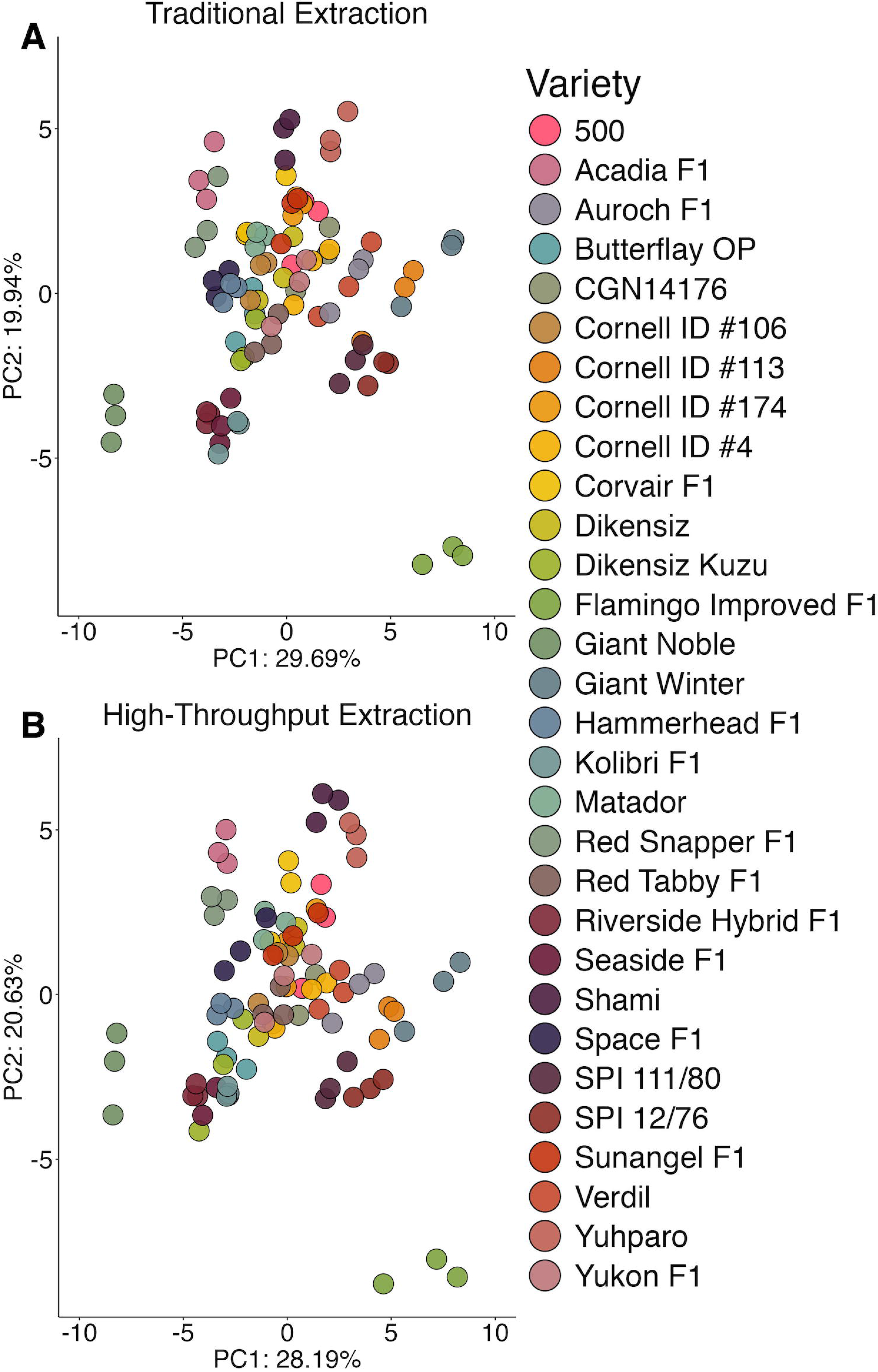
Principal components analysis 2-D scores plot displaying the population structure generated by the traditional extraction (A) or high-throughput extraction (B). Variable means were scaled to zero with a standard deviation of one.

To further explore small differences in population structures between extraction methods, we utilized random effects modeling to partition variance associated with genetic background, extraction method, replication, and within-method variability (Figure 3). In general, genetic background explained most variation (40.0% – 97.5%) compared to extraction method (0% – 49.0%). One notable exception is spinacetin (345 *m*/*z*) which was only detectable in our traditional extraction method. It suggests that spinacetin detected in the traditionally extracted samples was possibly from the hydrolyzed glycosylated flavonoids due to the low pH and long sonication times.

**Fig. 3.**
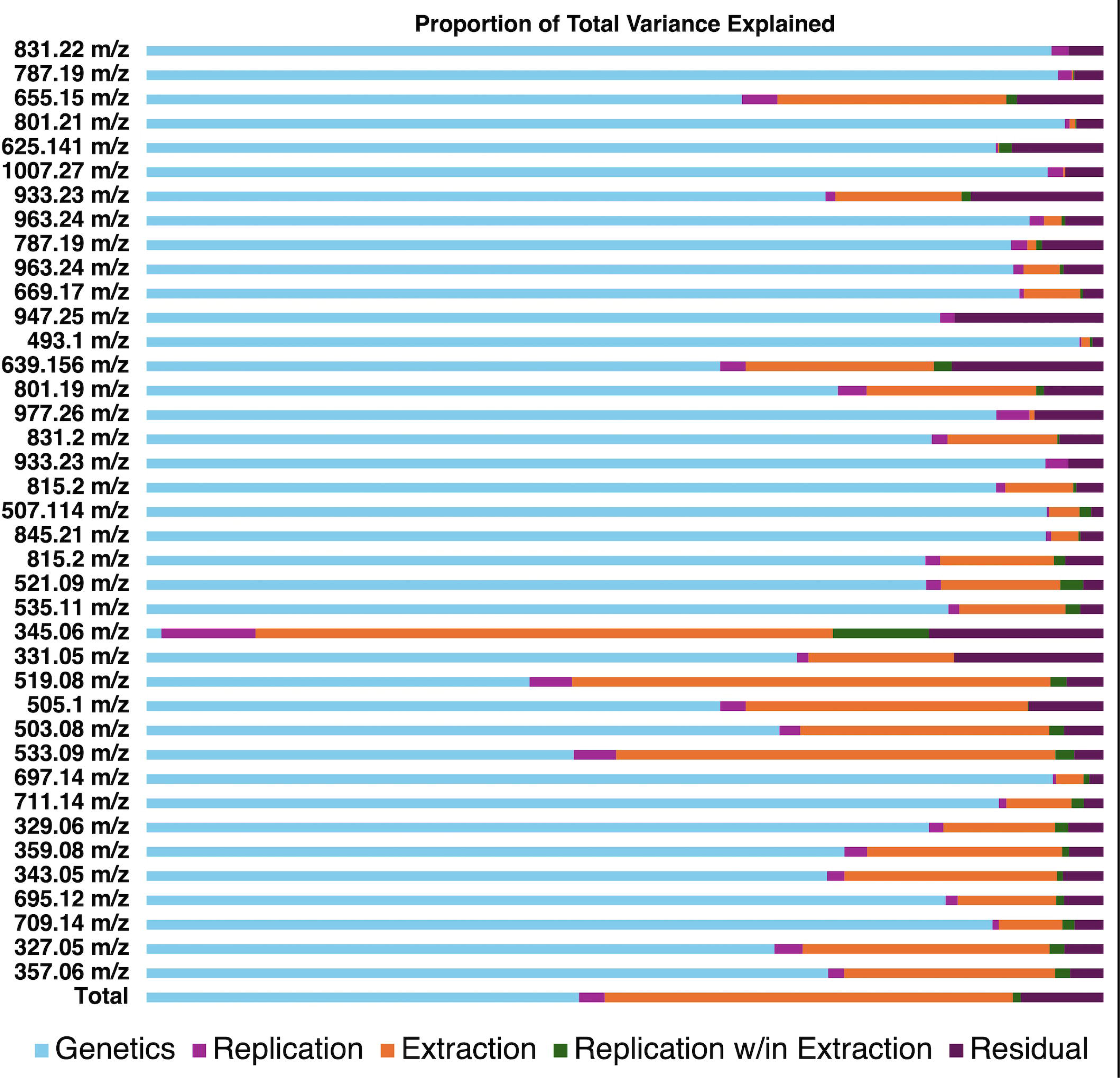
Visualization of variance estimates for each analyte profiled by our UHPLC-MS/MS method. Estimates were generated through random effects modeling separating the proportion of variance explained by genetics, replication, extraction method, replication within method, and residual error.

Another important trend is that the contribution to total variance caused by the extraction method tended to increase inversely to analyte polarity. Analytes in Figure 3 are in order of retention time from top to bottom which to a large degree is controlled by polarity. Analyte estimates were universally lower in the high-throughput extracted samples indicating that these later-eluting molecules were not extracted as effectively as with a sonication-based approach (Supplemental Table 3 and 4). Later eluting analytes in our method tend to be lower molecular weight. Given our observation about spinacetin, another possibility is that some molecules that are higher in the traditional extraction are hydrolysis products of larger, more glycosylated flavonoids. For example, 519.08 *m*/*z* could feasibly be derived from 695.12 *m*/*z* if the ferulic acid residue was hydrolyzed during extraction. However, there is no indication that pools of 695.12 *m*/*z* are being exhausted in the traditional extraction given that its estimated concentration is higher than that of the high-throughput method (Supplemental Table 3 and 4). Thus, differences between extraction efficiencies of these more nonpolar molecules is the most likely explanation. Previous work has shown that the breadth of polarities in spinach flavonoids requires different solvents depending on the analyte of interest (Singh et al., 2018a). Future work may merit exploring the use of different solvents or solvent mixtures at each extraction step to more broadly capture spinach flavonoids in the high-throughput approach.

A key advantage of the high-throughput method is that 48 samples can be simultaneously extracted in roughly 60 minutes while using significantly less solvent (3 mL/sample compared to 15 mL/sample). The high-throughput extraction approach could be utilized by a researcher interested in profiling a large population of spinach, and the traditional approach could be later utilized on a subset of samples that require additional accuracy and precision.

Here, we present the first comprehensive UHPLC-MS/MS method for spinach flavonoids as well as an accompanying high-throughput extraction method. Our method can separate and quantify 39 flavonoids uniquely produced by spinach in 11.5 minutes. Our UHPLC-MS/MS method features excellent sensitivity; low femtomole-on-column quantification limits enabling miniaturized extractions to be used in tandem. A limitation of our method is that the absence of authentic standards for spinach flavonoids make it impossible to know the true ionization efficiency of each molecule. Future studies are needed to establish ionization efficiency ratios between quercetin-3-glucoside and authentic spinach standards to more accurately estimate their concentrations. We also developed and tested a high-throughput extraction method (48 samples/hour; 3 mL solvent/sample), which can save researchers time and reduce solvent usage. The high-throughput extraction approach could be utilized by a researcher interested in profiling a large population of spinach, and the traditional approach could be later utilized on a subset of samples that require additional accuracy and precision for the more nonpolar flavonoids. Given sporadic reports in the literature, the data presented here represents benchmark values and ranges for many spinach flavonoids. Our tools enable these understudied analytes to be examined in a variety of contexts and open new research avenues in plant biology, food science, and human health research.

## ACKNOWLEDGEMENTS

We thank David Brenner and the USDA-ARS GRIN for providing some of the germplasm used in this study as well as Massimo Iorizzo of North Carolina State University for advice on curating this population. We also thank Brian Chlouber for assistance with plant growth and harvest. We appreciate the advice of Bhimanagouda Patil of Texas A&M University for suggestions regarding internal standards and extraction methodology.

## DISCLAIMER STATEMENT

The findings and conclusions in this publication are those of the authors and should not be construed to represent any official USDA or U.S. Government determination or policy. Mention of trade names or commercial products in this publication is solely for the purpose of providing specific information and does not imply recommendation or endorsement by the U.S. Department of Agriculture. The USDA is an equal opportunity provider and employer.

